# Curation at Scale with EPITOME: Extraction Pipeline for Immunological Texts and Open-Source Multimodal Enquiry

**DOI:** 10.64898/2026.01.08.698331

**Authors:** Eve Richardson, Parker Lischwe, Jason Bennett, Nina Blazeska, Jason Greenbaum, Blake Harlan, Noor Lallmamode, Daniel Marrama, Michael Talbott, Randi Vita, Alessandro Sette, Kaleb Kuether, Bjoern Peters

**Author notes:** co-first authors (Eve Richardson, Parker Lischwe).

## Abstract

The Immune Epitope Database (IEDB, iedb.org) has manually curated epitope data from over 26,000 publications across two decades. With PubMed adding ∼5,000 articles daily, traditional curation methods face scalability challenges. Given the multimodality of data contained in scientific papers, we have sought to build an open-source vision language model (VLM)-based tool that human curators can use to speed up and automate biological data curation. **Here we** present a multimodal document ingestion and Question-Answering (QnA) pipeline that ties traditional Optical Character Recognition (OCR) and text matching with Vision-Language Model (VLM) capabilities. The system, which we call EPITOME, implements three-stage processing: regex-based epitope and MHC molecule identification, visual element extraction from PDFs, and contextual indexing that links peptide sequences, MHC molecules, and assays to their locations across text, tables, and figures. This indexing is used to supply context for further VLM QnA. Our preliminary results from EPITOME demonstrate promising zero-shot performance of open-source VLMs that suggest promise for accelerating biocuration through a curator-in-the-loop process, with our evaluation identifying strategic points where curator-in-the-loop intervention can enhance overall system accuracy.

## 1. Introduction

Classical Major Histocompatibility Complex (MHC) molecules are a class of highly polymorphic cell surface protein complexes, whose function is to bind peptides (short amino acid sequences) and present them for recognition by T cells. Peptides that are recognized by T cells are referred to as epitopes (more specifically, T cell epitopes, with epitope being a general term to denote molecular structures recognized by the adaptive immune system)^1^. As MHC presentation is required for T cell recognition of a peptide, in-vitro assays measuring the binding of synthetic peptides to different MHC molecules (MHC binding assays) have long provided important insights into how immune recognition varies within a protein (identifying the most immunogenic regions of proteins), across variant proteins (in the context of infectious disease or cancer) and with respect to different MHC molecules (e.g. understanding how allelic polymorphism will affect immunogenicity of a peptide) ^2–5^.

Since its launch in 2004, the Immune Epitope Database (IEDB, iedb.org) has become the central repository for experimentally-validated B and T cell epitopes and MHC ligands across infectious disease, autoimmunity, allergy, and transplantation research domains^6–9^. The latest public metrics report 1.7 million epitope structures, supported by over 7.1 million assay records from over 26,000 references, all freely accessible through a web interface where users can build their data queries.

The IEDB team manually curates data from the literature into a structured format^10–14^. Common across the three main datatypes in the IEDB (B cell, T cell, and MHC assays), key properties describing experimental assays and epitopes must be codified from free text into a standardized format to enable querying. For each field for a given datatype, the curator inputs a value; for a subset of fields, this must be derived from a controlled vocabulary^15^. In practice, this is achieved in one of two ways: (i) data is entered via a web form, known as the Curation Application, or (ii) via a spreadsheet-like format, referred to as the Data Submission Tool (DST) template. Both methods convert unstructured data from text, figures, and tables present in scientific literature into the structured database format. The curation web form allows curators to enter epitopes and assays one at a time, while the DST allows curators to upload a large amount of data at one time. Given its critical role in processing large-scale datasets, herein we focus exclusively on DST-based curation.

The DST template’s tabular format contains a set of defined fields (columns) that describe in each row a specific observation (or assay) of an epitope-receptor interaction. For certain fields, there is a drop-down list that enables access to ontology terms; others (such as “Epitope Name”), are free text. The curator then submits the completed DST template to an in-house validation tool that checks for consistency between fields in each row, such as “Is the epitope sequence found at 100% identity within the sequence of its curated protein source?”. Issues are flagged by the validation tool and the curator will refine their initial inputs until all validation checks are passed.

Given the labor intensiveness of the IEDB curation process, we have continuously tried to automate aspects of it by leveraging approaches developed by the Natural Language Processing (NLP) community^4^. However, the application of NLP tools that have shown good performance in general information retrieval tasks to immunological data curation has, in our experience, been challenging and not trivial. One recurring challenge is that much of the curatable information is embedded in visual elements (e.g., figures and tables) rather than the main text of publications. While optical character recognition (OCR) enables access to some of the information contained within these visual elements, this still leaves more complex textual data (e.g. with variable spacing or where the order of elements must be understood with reference to an image) and non-textual data (such as graphs) inaccessible.

Furthermore, while there are several NLP methods trained specifically on biomedical literature that achieve high performance on subtasks within our curation process (specifically, entity recognition and relationship extraction tasks such as gene symbol recognition), linking these extracted entities and relations into our specific data format has not been straightforward. Perhaps one of the most significant challenges to automating curation is the large number of synonyms, variants, and context-dependent meanings that are used not only by different researchers, but even within a single paper. While a curator reading the paper can make inferences, codifying these inferences into rule-based systems would be extremely challenging, requiring extensive manual feature engineering for each new variation.

The most significant challenge is our requirement for complete curation of a single document with very low error tolerance. This differs significantly from most NLP/LLM tools, which excel at extracting and summarizing information from thousands of documents at speeds no human can match, but generally accept some level of error as a trade-off for scale and speed of responses. This approach is incompatible with the IEDB’s requirement for accurate capture of all relevant data points from each publication: we are entrusted with accurately capturing every epitope, every assay condition, and every immune response measurement, with an error rate of more than 1% being unacceptable. As a result, such tools will require significant safeguards against the introduction of errors. We therefore envisage their use only in a “curator-in-the-loop” scenario in which our highly-trained curators will validate or correct inputs prior to validation.

In 2024, the average curation rate amounted to 0.9 days / publication / curator, however this is highly non-uniform: an increasing number of longer papers (reporting a larger number of distinct epitopes and assays) can take several days to curate via the DST route. We estimate that around 50% of this is required for data entry alone with an additional 20% for subsequent validation and correction rounds (and the remainder required for reading the publication). It is with these time-consuming papers that we anticipate the most potential for automation (rapidly performing data entry from large tables) and the most significant gains from automation.

In late 2024, the IEDB entered a collaboration led by Intel Corporation to showcase how LLM technology can benefit biomedical science. LLMs are deep learning models trained on vast amounts of natural language data. One of their main advantages is their success on tasks on which they have not been explicitly trained (zero-shot settings), with significant improvements upon this strong baseline available by providing a limited number of labelled examples (few-shot learning; specifically in LLMs, this does not entail finetuning but rather supplying these examples at inference time)^16,17^. This means that LLMs do not require the large corpuses of annotated data required by traditional NLP methods, meeting or exceeding performance of fully fine-tuned models with thoughtful use on several literature examples of biomedical NLP tasks^18–20^. At present, multiple open-source large language models exist, such as LLaMA, Mistral, Gemma and Qwen, which can be used for custom development and fine-tuning^21–24^.

Recent advances have expanded the capabilities of LLMs into the visual domain, allowing graphical information to be processed by prepending a visual encoder that maps visual embeddings into a shared latent space for the LLM to use as context before generating a response. These visual language models (VLMs) are trained end-to-end on image-text pairs, generating text just like a traditional LLM, but are able to describe or answer queries pertaining to a visual input. The advent of these VLMs therefore now enables direct query of visual data elements within scientific papers^8^. Current open-source VLM models include Large Language and Vision Assistant (LLaVA) and Qwen-VL^24–26^.

While VLMs can directly use images (such as full pages of a paper) during query, supplying full papers during prompting raises issues with context window length; furthermore, it has been shown that supplying the LLM with irrelevant context may significantly reduce response accuracy, and that response accuracy declines when the relevant information is in the middle of large context window, as is likely to be the case for scientific papers^27,28^. Furthermore, supplying more granular context to the VLM improves interpretability. Modern document processing frameworks, like Docling, enable this, transforming article PDFs into structured formats that can be used to generate more precise contexts to use for VLM query ^29^.

## 2. Methods

### 2.1 MHC-Bind dataset

We created a dataset from a subset of papers reporting MHC binding assays which we refer to as *MHC-Bind*. MHC binding assays were selected as a starting point due to their relatively simple nature as compared to B and T cell assays. The IEDB was queried to return a list of publications containing only MHC binding assays, with PMCIDs. This set was further limited to those publications where the epitope linear peptide sequences were described in the main sections of the publication, rather than in supplemental figures or tables. Non-peptidic epitopes and epitopes with modifications were further excluded.

### 2.2 EPITOME Document parsing

Our document parsing pipeline begins with document ingestion via Docling using default *tesseract* OCR parameters^29,30^. An additional OCR step was applied to extract text from images using *pytesseract*^31^. We supplement Docling’s native methods for pairing text captions (headers and footers) and visual elements with custom logic for finding the nearest text indicating a Table or Figure header and footer. The output of the document parsing step is the full text, and each visual element is paired with the corresponding header and footer text.

### 2.3 EPITOME Identification and standardization of entities

To identify both peptide linear sequences (which are sequences of consecutive valid amino acid characters) and MHC protein complex names (which generally conform to species-specific nomenclature rules, with established synonyms as per the MHC Restriction Ontology), regular expression matching is first used on text extracted from the main text, tables and images^32^. To further improve recall of values which are intractable to OCR, images and tables are directly supplied to the VLM, seeking identification of linear epitope sequences and MHC protein complex names.

To identify Assays, we query each visual element to identify the Assay Response(s) and Units. For valid Assay Response(s), we then re-query the visual element with a restricted list of possible Assay Types specific to the predicted Assay Response alongside their full textual descriptions as per OBI^33^.

For validation, MHC protein complex names were standardized with a custom function using synonyms as previously compiled in the MRO^32^. Peptides were filtered for valid amino acid strings of length at least equal to 3. Putative peptide matches were further filtered using English stop words, which were compiled using *nltk.corpus.stopwords*^25^*, sklearn.feature_extraction.text.ENGLISH_STOP_WORDS* and *wordfreq’s* top 5e-4 English words^34–36^. Assay Type and Response validity was enforced via restricting responses to valid OBI names.

### 2.4 EPITOME QnA and agentic calling

For MHC Allele Name and Source Organism Taxonomic ID, the number of recognized synonyms within the respective ontologies is too large to fit within the context (90,187 MHC synonyms and 3,501,885 organism name synonyms in the IEDB’s organism tree). Custom parsing functions were used to extract and standardize the relevant values using the Aho-Corasick algorithm (*pyahocorasick*) and NCBI’s *Entrez* taxonomy utilities^37^.

For the Source Antigen field, Qwen’s agentic behavior is used to run PEPMatch where the Source Organism corresponds to a valid proteome^38^.

### 2.5 VLM implementation

The Vision Language Model we deploy is Qwen2.5-VL-32B-Instruct as it is designed to excel at objective queries such as logical reasoning and knowledge-based Q&A, delivering enhanced detail and clearer formatting for critical use cases^24^. This model is also made available through vLLM Habana-fork for continuous inferencing on Intel’s Gaudi AI accelerators, which we use for high throughput and efficient resource utilization. We explored utilizing a small locally hosted model that can be deployed on common consumer grade hardware especially with quantization, choosing the 32B Qwen2.5 vision language model at FP16 precision. The model was served via Habana AI’s vLLM fork on a single 2x Xeon 8x Gaudi 2 server. The Qwen2.5-VL models have a 128k context window, allowing for all relevant context to be included within a single query. Model confidence for a given response was calculated as the average log probability calculated across all output tokens.

### 2.6 Evaluation metrics

We describe performance via sensitivity (%), precision (%), calculated versus corresponding validated human curator-reported values. For Source Organism ID, we bin predictions by the level of their taxonomic match: Exact Match, More Specific Taxonomic ID (where the prediction is a child of the true taxonomic ID), Less Specific Taxonomic ID (where the prediction is a parent of the true taxonomic ID), Incorrect (where there is no relationship between predicted and true taxonomic ID), and Unknown (where the VLM returns None).

## 3. Results

### 3.1 MHC-Bind: an annotated dataset of MHC binding data

For the development of our VLM curation prototype, we created a dataset from papers containing MHC binding data that have already been curated in the IEDB. We report data from a set of 251 publications reporting 6,418 peptide-MHC measurements. This is referred to as “MHC-Bind”.

MHC binding assays are in-vitro assays that measure the binding of synthetic peptides and MHC molecules. The interaction of a given MHC molecule and peptide are measured using one of several assay formats (**Table S1**). Each row in MHC-Bind corresponds to an observation of a peptide-MHC molecule interaction within a paper. As each paper may contain multiple peptides, multiple MHC molecules and multiple assays, successful curation involves not only accurat identification of these subcomponents (peptide, MHC molecule, and assay) but linking in a many-to-many-to-many format (**Figure 1**).

**Figure 1:**
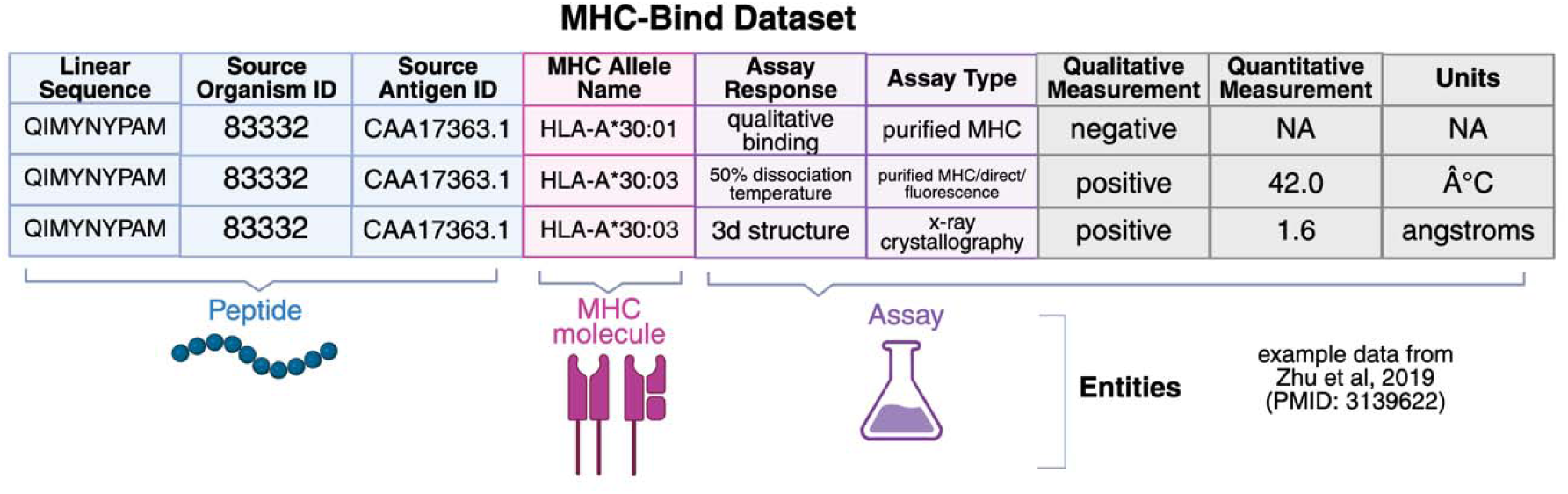
MHC-Bind is a tabular dataset of observations of peptide-MHC assays, where each row corresponds to a **Peptide-MHC protein complex** interaction measured via an **Assay.** Peptides are identified via their **Epitop Linear Sequence**, and have the corresponding properties of the **Source Antigen ID** and **Source Organism ID**. Th **MHC protein complex** has the property of the **MHC Allele Name** which adopts a value from the MHC Restrictio Ontology (the IEDB name of the corresponding CURIE)^22^. Assays are described via the **Assay Response** the measure and the **Assay Type**, which adopt values from the Ontology of Biomedical Investigations^21^. Each peptide-MHC measurement has an associated qualitative and, for non-qualitative assays quantitative, measurement with a associated unit. Qualitative Measurement, Quantitative Measurement and Units are marked here in gray as we d not tackle these fields in the present work.

We selected a subset of tractable columns related to each of these three key elements from the full DST used by curators (**Figure 1**), described in **Table 1**. With these entities and fields in mind, we devised a strategy for automated curation.

**Table 1:**
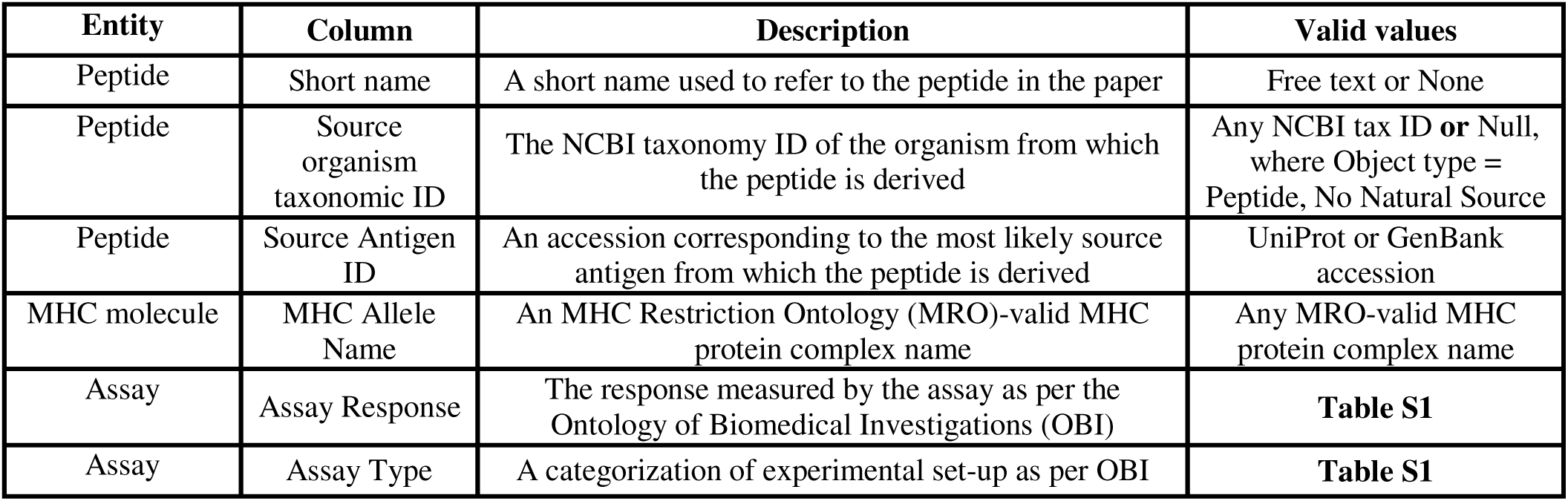
We targeted several columns in the DST field, each related to one of the three entity types (Peptide, MHC molecule, Assay). The additional fields of Epitope Object Type and Epitope Structure Definition are defined in **Table S2**.

### 3.2 EPITOME: a VLM-based approach for partial automation

We devised an end-to-end pipeline starting from a manuscript PDF and culminating in a partially-completed DST as illustrated in **Figure 2**. The process uses VLMs for several steps: to improve sensitivity of peptide- and MHC allele name extraction (**Figure 2B**), to enable recognition of visual elements in which Assays are performed (**Figure 2B**), to complete chosen DST fields (**2C, 2D**) via QnA and agentic tool calling, and to pair up peptides, MHC molecules and Assays within visual elements (**Figure 2E**). The result is then used to assemble the DST template. In practice, this is intended as a “curator-in-the-loop” process. The curator must inspect the VLM-supplied outputs: these can be removed, accepted or corrected, or the query can be repeated altogether. In **Figure 2**, the person icon indicates where the curator’s input is required.

**Figure 2.**
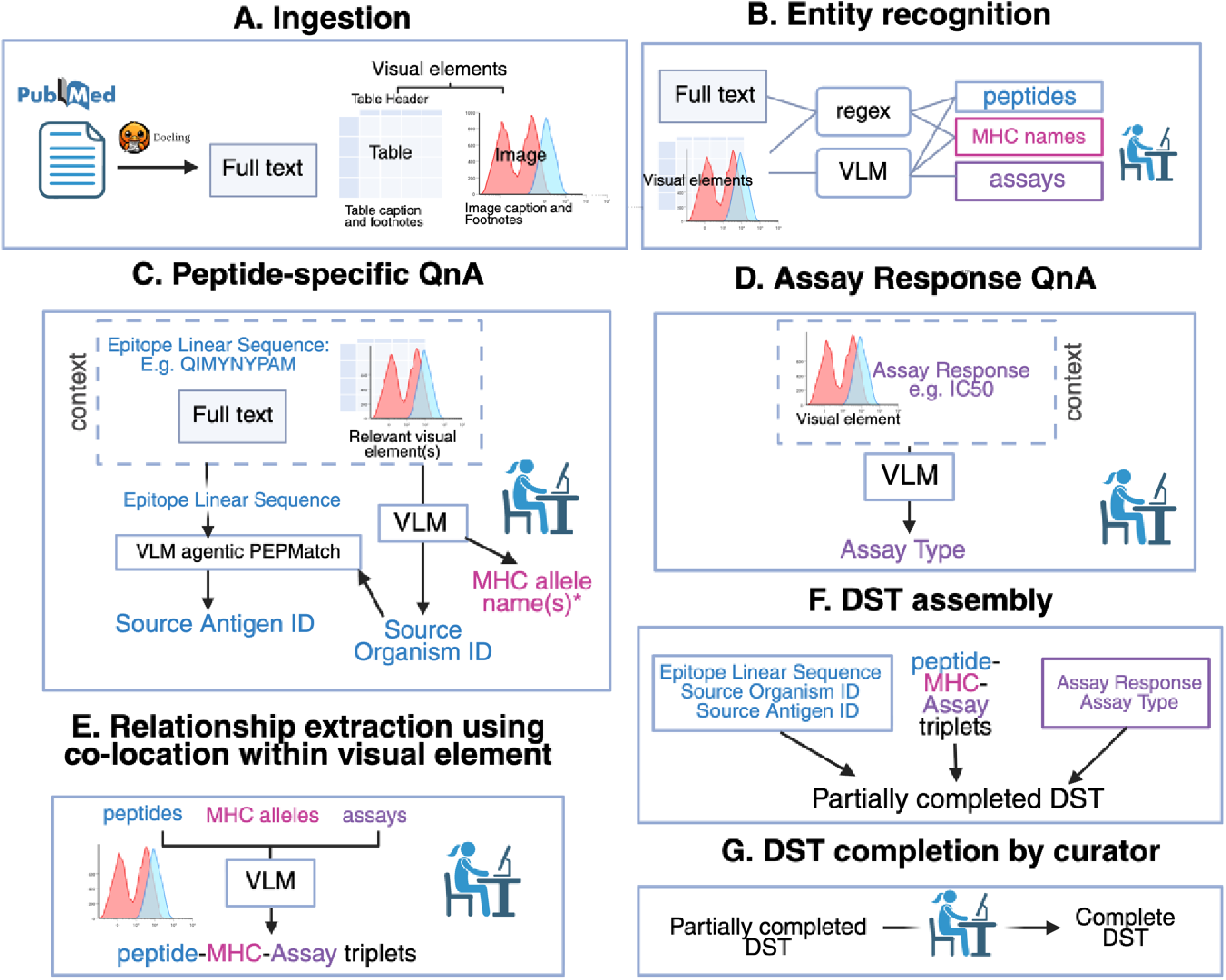
The first step involves ingestion of the PDF into its full text and visual elements (**A**). The text and visual elements are used as input for recognition of the three key entities in MHC-Bind: peptides, MHC allele names an Assays. For peptides and MHC allele names, this is achieved via a combination of regex and directly querying VLMs with each visual element. For Assays, each visual element is interrogated for any Assay Responses measure and reported within the figure (**B**). For each unique peptide, the full text and relevant visual elements in which it appears are used as context for VLM query to answer peptide-specific fields such as the Source Organism ID. Source Organism ID and the peptide’s sequence are used by the VLM to return a Source Antigen ID. The VLM is also queried to produce a peptide-specific list of MHC alleles, i.e. a list of MHC alleles against which a specific peptide is tested. (**C**). For each visual element in which an Assay Response is putatively measured, the VLM is queried to return the appropriate Assay Type given the Assay Response (**D**). Each of the three entity types identified within the same visual element (as well as the peptide-specific list of MHC alleles or the paper-level MHC allele list, if no MHC molecules are co-located) are supplied to the VLM for pairing into triplets (i.e. pairing into which peptides were tested with which MHC molecules and via which assay) (**E**). In **G**, EPITOME uses each of the fields calculated in **C** and **D** and the mapping from **E** to assemble a partially-completed DST, which is complete manually by the curator (**F**). The icon of the person with the computer represents the “curator-in-the-loop”, and is included where the curator must inspect VLM-supplied values, either confirming, correcting or re-querying.

In designing EPITOME as a curator-in-the-loop tool, we considered two main factors, which were minimum acceptable performance (maximal sensitivity with a minimum acceptable precision) and maximizing interpretability and flexibility for the curator. We aim to have maximal sensitivity for each form of entity extraction. However, we do not want the curator to be inundated with false positives (e.g. incorrect peptides, MHC allele names or Assays): a precision of 1%, for example, would require the curator to discern 99 false positives for every true positive. We select a minimum acceptable precision of 10%. Interpretability and flexibility is increased greatly (over a simple baseline of supplying full paper PDFs), by a multi-step process with multiple points of intervention (**Figure 2**) for correction or re-prompting. Reducing the context supplied to the VLM means that the curator can also inspect and correct this, e.g. by adding additional relevant visual elements during the peptide-specific QnA.

Ultimately, EPITOME will be served as a graphical user interface (GUI) with intervention at each step. For the following methods, however, we explore zero-shot performance at each of its component steps (**Figure 2B, C, D, E**).

### 3.3 VLMs enable zero-shot recognition of key entities in 70% of papers

The first step in EPITOME is to identify mentions of peptides. This is achieved via a combination of regular expression matching against text extracted via OCR and VLM queries for visual elements. Inspection of visual elements is necessary given 95.2% of epitope sequences and 99.6% of assays are reported to lie in figures and tables in MHC-Bind (**Figure 3A**).

**Figure 3:**
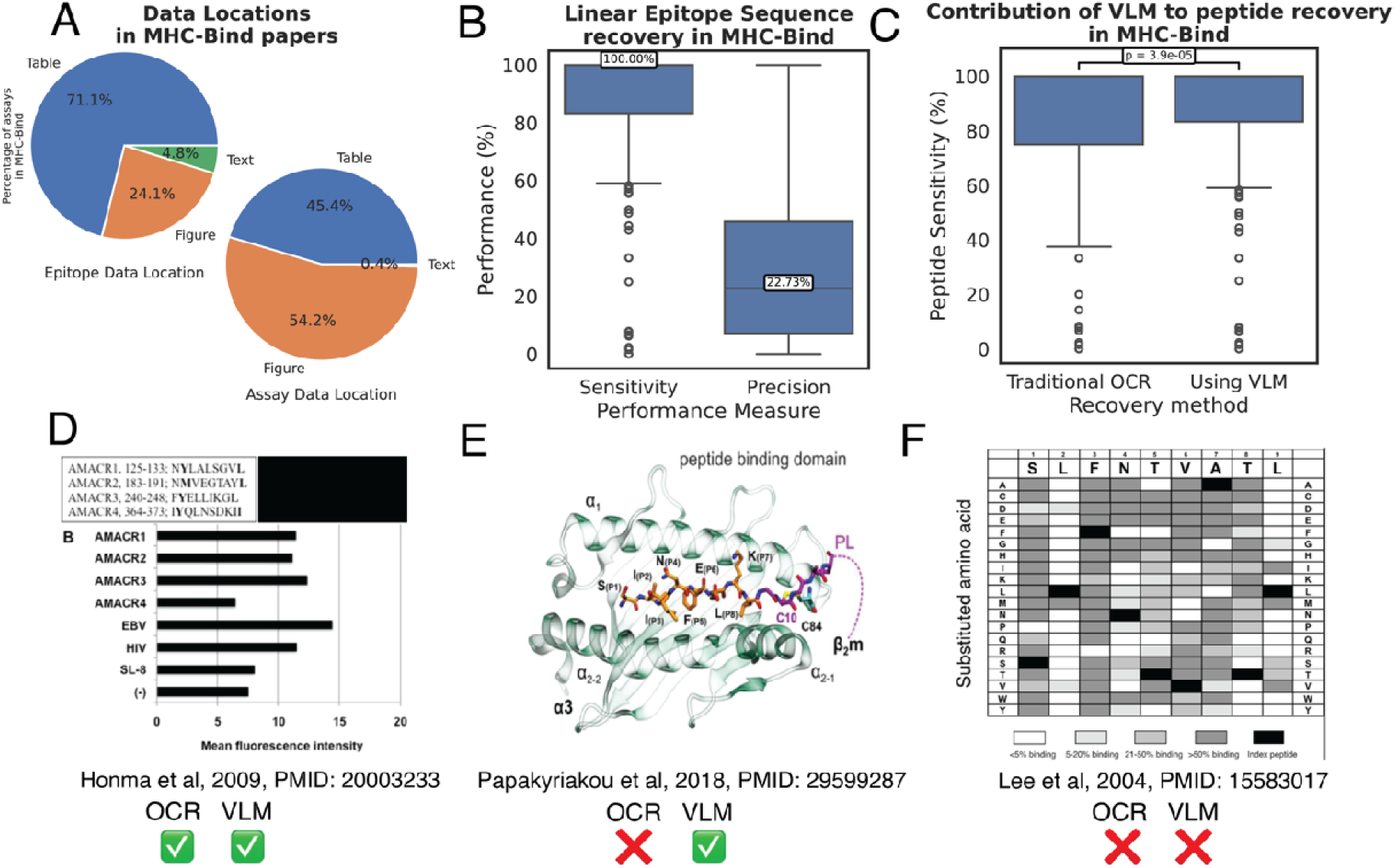
Given that 95.2% of peptides sequences and 99.6% of assays in MHC-Bind are reported in visual elements (**A**), EPITOME’s ability to extract data from visual elements is critical. EPITOME extracts relevant data from visual elements using both OCR and direct VLM query, with a median sensitivity of 100% and precision of 22.73% (**B**). The VLM makes a significant contribution to sensitivity (**C**). For figures such as that reproduced from Honma et al (2009) in panel **D**, the peptide sequences are represented in plain text and can be extracted using OCR^39^. The principal advantage of VLMs is their ability to parse data in abstract formats, such as labelled molecular diagrams like this pMHC molecular visualization from Papakyriakou et al (2018)^40^ (**C**). Nonetheless, there remai challenging examples which Qwen2.5-VL models (both 32B and 72B) were unable to parse, for example this combinatorial representation from Lee et al (2004)^5^ (**D**). In this example, single substitutions at each position of a length 9 peptide (SLFNTVATL) were tested, resulting in a total of 19 X 9 = 171 peptides. The VLM was able t extract the reference peptide, but not the 171 derivatives for which data is presented in the figure. Figures are reproduced from their respective publications under a CC-BY-2.0 license (**C**), CC-BY-4.0 license (**D**) or CC BY-NC-SA 4.0 license (**E**).

We demonstrate that EPITOME achieves 100% sensitivity for 70.5% of papers in MHC-Bind. In this first pass without curator intervention, EPITOME achieves a median precision value of 22.7% (**Figure 3B**). We note that 20.1% of false positives are exact matches to peptides already curated in the IEDB (i.e., they are mentions of peptides from real proteins that have been tested elsewhere in MHC or T cell assays, but are not relevant to the specific MHC binding assay being curated). A further 18.5% of false positives have a Levenshtein ratio of ≤ 0.1, suggesting possible OCR error.

While most linear peptide sequences are reported in an OCR-tractable format, using VLMs enables a significant increase in sensitivity compared to traditional OCR alone (68.9% of papers with perfect sensitivity vs. 70.5%; p = 3.9e-5, Wilcoxon paired test, **Figure 3C**). **Figure 3D** shows an example of a peptide sequence representation which allows peptide extraction via both OCR and VLM query. **Figures 3C** and **3D** highlight two examples where the epitope linear sequence cannot be extracted via OCR alone but may (**3C)** or may not (**3D**) be amenable to the VLM approach. The situation portrayed in **Figure 3D** is particularly challenging, as it requires the curator to enumerate the displayed peptides from the heatmap, with exhaustive single position substitutions at each position of a length 9 reference peptide resulting in a total of 171 derivative peptides, in addition to the reference peptide itself.

After the extraction of peptides from papers, EPITOME uses the MHC Restriction Ontology (MRO) to robustly identify MHC protein complex names via a similar combination of regex and VLM query. This achieves comparable sensitivity (100% sensitivity for 68.5% of papers) with a median precision of 16.7%.

Unlike linear epitope sequences or MHC molecule names which can be extracted using simple pattern matching on OCR-extracted text, the third component of MHC-Bind, the Assay object, requires more sophisticated methods for extraction. As per **Figure 2**, we directly queried each visual element for Assay Response using the VLM, with a chained query to identify Assay Type given the Assay Response extracted. On a per-paper basis, using the VLM to this end achieved an average sensitivity of 78.6% and average precision of 45.6% for recognition of Assay Response, achieving 100% sensitivity for 73.7% of papers. For recognition of Assay Type, average sensitivity was reduced to a sensitivity of 35.1% and precision of 14.5%, achieving the desired 100% sensitivity for 28.5% of papers.

### 3.4 The RISEN framework for query construction enables the zero-shot use of a VLM for successful QnA

Subsequent to entity recognition, EPITOME’s peptide-specific context is supplied to the VLM for the QnA procedure used to fill in the peptide-specific fields of the DST.

We construct queries that include these multimodal elements without embedding retrieval in the pipeline. Multimodal queries are developed for each extracted epitope by routing elements from the epitope’s known locations to the user prompt. We follow the **RISEN framework** when constructing queries to improve accuracy. We do so by first specifying the **R**ole of the VLM through the system prompt, addressing what the model should focus on: “You are an intelligent AI assistant that is an expert in understanding and answering questions about immunology. As an immunology expert, you are given snippets from real immunology papers, and you are tasked to extract specific information from the text and images provided as context.” Once the role is defined, we create the user query which contains the Instruction, Steps, Expectation, and Narrowing targeted for the corresponding DST column we are curating. The **I**nstruction contains the simple command for what to answer, and we contextualize this command by supplying textual information from multiple sources already available to actual curators, including **S**teps as information from the IEDB curation manual which are used to ensure consistency and accuracy of literature-based curation^41^.

Beyond the domain-specific information supplied by the curation manual, we also provide the acceptable outputs, where possible given context length, for DST columns to **N**arrow the possible outcomes generated by the model. Finally, we specify the **E**xpected output in terms of formatting, which includes an example output that adheres to the expected format of the corresponding DST column entry.

We used this RISEN QnA framework to answer the “Source Organism ID” field (**Table 1**). Responses were evaluated for correctly identified peptide sequences from the PMID set (2,227 peptides from 235 papers). EPITOME returned an exact match for 73.4% of values. For a further 15.5% of values, EPITOME returned a taxonomic identifier in the correct lineage of the tree, but with either a less specific (14.2%) or more specific (1.3%) taxonomic identifier than was curated. For example, an epitope was curated as originating from ‘Influenza A virus’, but EPITOME returned a the taxonomic child ‘Influenza A virus (A/Puerto Rico/8/1934(H1N1))’ (**Figure 4A**). Another promising result is that model confidence was significantly higher for correct taxonomic identifiers than incorrect, meaning that model confidence could be a useful tool for curators in scrutinizing VLM output (**Figure 4B**). We also explored performance on two other fields, demonstrating again that model confidence can be used to separate correct and incorrect predictions, with ROC-AUCs of 0.81 and 0.91 (**Figure S1**).

**Figure 4:**
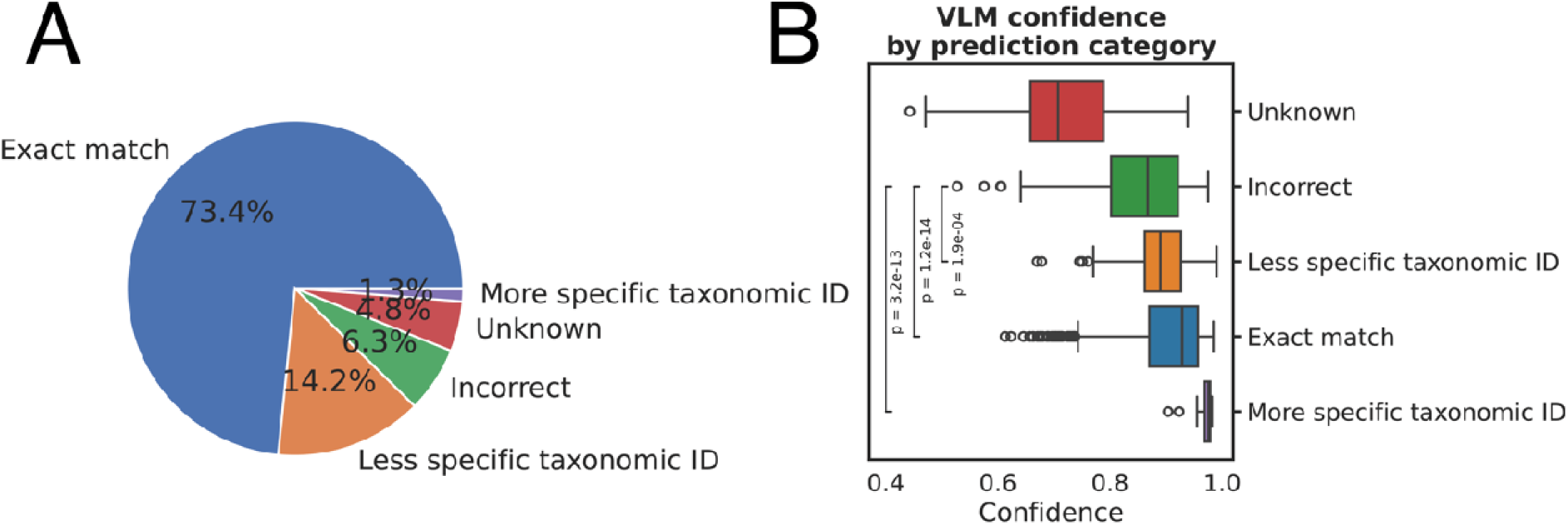
With EPITOME, we assessed zero-shot performance of Qwen2.5-VL-32B in QnA for the completion of peptide-specific fields in the DST. We show results here for the Source Organism Taxonomy ID. EPITOME achieved an exact match to the ground truth for 73.4% of entries and returned a taxonomically related suggestion for an additional 15.5% of entries **(B**). Examining VLM model confidence (**C**) showed that incorrect taxonomic suggestions (6.3%) were significantly less confident than less specific, more specific or exact taxonomic matches (values calculated via Mann-Whitney-U).

### 3.5 Extending VLM capabilities with agentic tool calling

Although most curatable information is contained within the papers themselves, certain fields require curators to query external databases, for example “Source Antigen ID”. “Source Antigen ID” is an identifier for the protein that the peptide is derived from in the Source Organism’ proteome. Curators use PEPMatch to search for peptides in the target proteome: where a match i found, PEPMatch returns a protein identifier (either from UniProt or the National Center for Biotechnology Information (NCBI) protein databases) which directly populates “Source Antigen ID”^38,42–44^.

To integrate this into the automated curation workflow, we launch our VLM with tool calling enabled, and access to a helper function that makes PEPMatch calls using the Next-Generation IEDB Tools API^45^. We used the VLM-assigned taxonomic ID to identify the appropriate proteome from the list of proteomes available in the PEPMatch API.

### 3.6 Evaluating the full EPITOME output

EPITOME’s combined output thus far is, for a given paper, a list of possible peptide sequence and associated fields, MHC molecule names and Assays. Each of these candidate lists can be traced back to visual elements. as shown in **Figure 5** with an example reproduced from Felieu et al, 2013.

**Figure 5:**
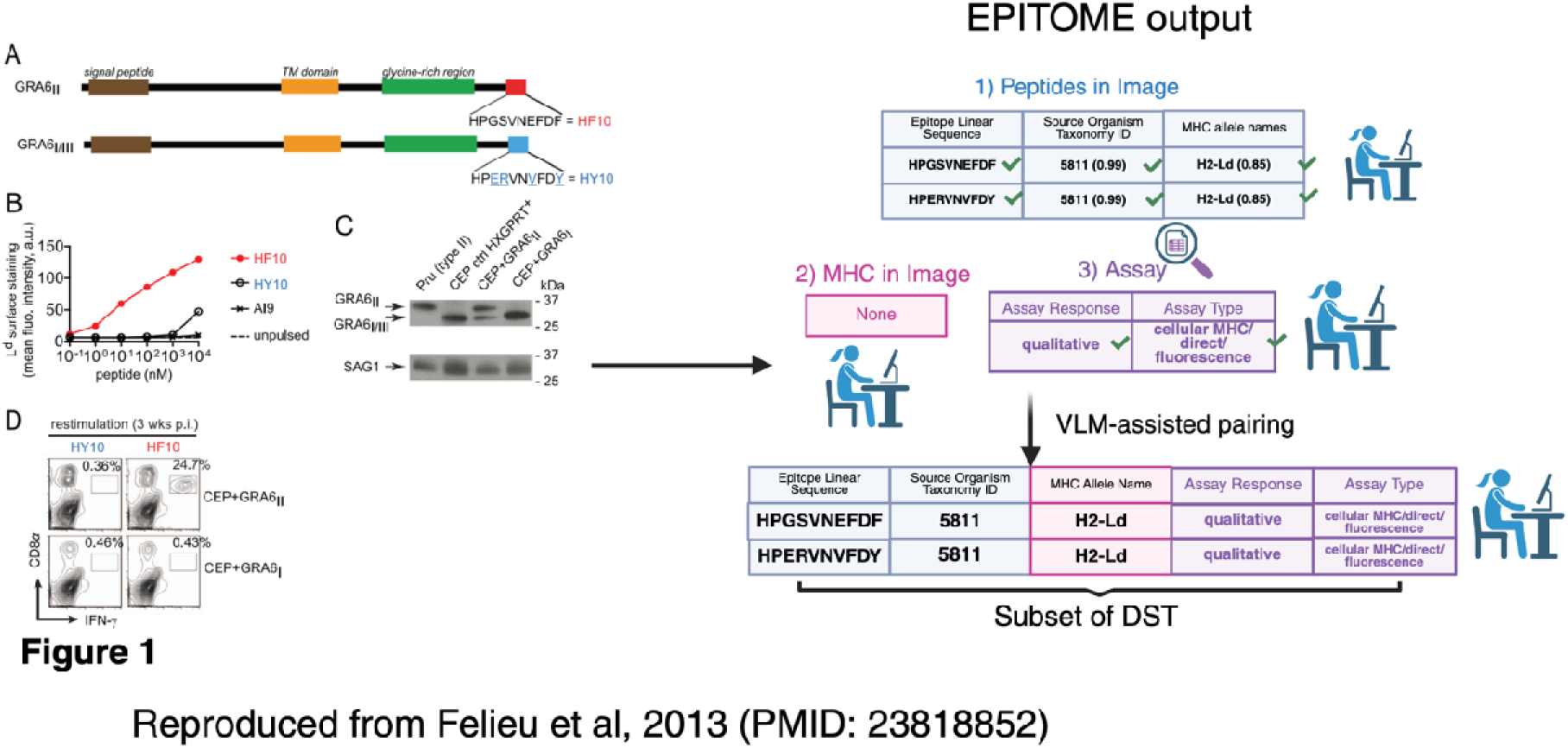
An example of EPITOME’s outputs for a given visual element in a paper (Felieu et al, 2013; PMID: 223818852). This is an example in which EPITOME correctly extracted all peptide, MHC and Assay triplets from the curated paper with 100 % sensitivity and 100% precision.

The final challenge is to pair each of the elements into valid rows in the DST (with triplets of MHC allele names, Assays, and Peptides; **Figure 5**). We explored whether this could be achieved using the co-localization of these entities within visual elements, using a VLM query to make the mapping. Using only correctly-identified peptides and associated assays, pairing peptides and MHC molecules was achieved with 100% sensitivity for 39.6% of papers. There was a similar success rate for pairing peptides and assay responses (37.7%). Due to the difficulty of correctly predicting Assay Type, just 11.3% of papers achieved 100% sensitivity in combining peptides with the correct Assay Type and Response. EPITOME identified all reported Peptide-MHC-Assay triplets in 8.3% of papers.

## 4. Discussion

The data in IEDB is the culmination of more than 20 years of manual curation parallel with community ontology development efforts^1,3,30^. This has resulted in the distillation of B and T cell epitope and MHC ligand assays from over 26,000 immunology publications (as of November 2025) into a highly standardized format adhering to FAIR principles^15^. The complexity of the curation of these papers, necessitating full-time PhD-level curators, means that integrating open-source VLMs into curation constitutes a true test of these technologies; meanwhile, our existing efforts towards machine-actionable interoperability and large corpus of richly annotated data mean that we are primed to take on the challenge.

EPITOME is our first step towards integration of LLMs into the curation pipeline. In this initial development phase, we have focused on a single datatype (MHC binding assays) creating a dataset that we refer to as *MHC-Bind*. In MHC binding assays, the interaction of a given MHC molecule and peptide are measured using one of several assay formats. The task requires successful identification of these key entities as well as answering questions about each. VLMs were essential to our process: they enabled a significant improvement in sensitivity over traditional methods alone for recognition of peptides and MHC molecule names, and enabled us to tackle a third entity type, the Assay object, without any finetuning, recognizing 100% of the assay responses measured in nearly ¾ of papers. They also demonstrated promise for completing DST fields, with greater than 73.4% exact match accuracy and 88.9% accuracy allowing for less or more specific taxonomic identifiers, with correct predictions having significantly higher model confidences than incorrect predictions. The clearest deficit in performance was in the relatively poor zero-shot performance of Assay Type classification, suggesting that either fine-tuning, further prompt optimization, or superior models are required for this to reach acceptable accuracy.

Our prospective use of EPITOME will require multiple safeguards against hallucination. An inherent limitation with today’s open-source LLMs is their ability to produce hallucinatory behavior, which are responses that contain convincing and confident language, while being factually incorrect or not pertaining to the input context. One of the ways we combat model hallucinations is by constructing prompts that limit the amount of context provided from the paper, as well as providing context that is grounded in highly confident, often deterministic content. Nonetheless, hallucinations remain an issue: for example, when extracting assay responses, we supply individual tables and figures extracted from the article and provide a controlled vocabulary of possible responses to make model output more deterministic. Nevertheless, there are still instances where the model gives an answer that is pertinent to the supplied visual information but does not follow the controlled vocabulary (**Figure S2**).

A curator-in-the-loop scenario is the ultimate protection against the possibility of inserting hallucinatory content into the database and will lead to substantial improvements in accuracy versus values reported here, without interim correction for chained inferences. Our next step is to integrate EPITOME into a graphical user interface (GUI), where the curator is able to correct or remove inferences as well as re-prompt. Through curator testing, we will be able to quantify EPITOME’s effect on curation rate.

Compared to the leading commercial models that range in hundreds of billions to trillions of parameters, we opted to use a small VLM (32B) that can be targeted for inference on consumer grade hardware. We saw minimal improvement using Qwen-VL-72B vs. our default, 32B (data not shown). For further improvements, we would look to a reasoning model to see whether zero-shot performance on the most complex tasks, such as assay type classification, can be improved without changes to context or prompting. There are few VLM reasoning models; prior to 2025, the only open-source multimodal reasoning model was QVQ-72B-Preview, a research model released by Qwen. Since then, there have been multiple other candidate models released such as Google’s gemma series, and Moonshot’s Kimi-VL^23,46,47^.

As we scale the solution to address a wider range of papers, and more templates that include different fields, we also look to integrate more rule-based tools (as we did with agentic calling of PEPMatch) and additional ontologies within the OBO Foundry^31^. We currently use ontologies only within the standardization of VLM outputs; one exciting avenue would be the development of agentic methods that allow the VLM to directly use ontologies at the time of inference.

Our VLM-based curation automation tool has shown promising performance on this highly specialized task that is only entrusted to PhD-level, trained curators. Continued testing from curators and interaction through a GUI will be the next steps to refine EPITOME’s approach and establish a robust curator-in-the-loop framework.

## Acknowledgements

We thank Intel Corporation and the La Jolla Institute for Immunology for computational and personnel resources, and the IEDB curation team for evaluation assistance.

