## Supplementary Materials for "Curation at Scale with EPITOME: Extraction Pipeline for Immunological Texts and Open-Source Multimodal Enquiry"

### **Supplementary Material**

### **Curation at Scale with EPITOME: E**xtraction **P**ipeline for **I**mmunological **T**exts and **O**pen-Source **M**ultimodal **E**nquiry

#### Eve Richardson^1#^, Parker Lischwe^2#^, Jason Bennett^1^, Nina Blazeska^1^, Jason Greenbaum^1^, Blake Harlan^2^, Noor Lallmamode^2^, Daniel Marrama^1^, Michael Talbott^1^, Randi Vita^1^, Alessandro Sette^1,3^, Kaleb Kuether^2*^, Bjoern Peters^1,3*^

#### * Corresponding author

#### # Co-first authors

#### Affiliations

#### ^1^ La Jolla Institute for Immunology, La Jolla, CA, USA

#### ^2^ Intel Corporation, Santa Clara, CA, USA

#### ^3^ Department of Medicine, University of California San Diego, La Jolla, CA, USA.

| **Assay Response** | **Allowed Assay Types** |
| --- | --- |
| **3D structure** | x-ray crystallography,  electron microscopy |
| **50% dissociation temperature** | purified MHC/direct/fluorescence |
| **MHC binding** | High throughput multiplexed assay |
| **association constant KA** | cellular MHC/direct/fluorescence,  binding assay |
| **binding constant** | binding assay |
| **dissociation constant KD** | purified MHC/competitive/fluorescence,  cellular MHC/competitive/fluorescence,  cellular MHC/competitive/radioactivity,  lysate MHC/competitive/radioactivity,  purified MHC/competitive/radioactivity,  binding assay |
| **dissociation constant KD (~EC50)** | purified MHC/direct/fluorescence |
| **dissociation constant KD (~IC50)** | purified MHC/competitive/radioactivity,  purified MHC/competitive/fluorescence |
| **half life** | purified MHC/direct/radioactivity,  cellular MHC/direct/fluorescence,  purified MHC/direct/fluorescence,  lysate MHC/direct/radioactivity,  cellular MHC/direct/radioactivity,  binding assay |
| **half maximal effective concentration (EC50)** | lysate MHC/direct/radioactivity,  cellular MHC/direct/fluorescence,  purified MHC/direct/fluorescence,  binding assay |
| **half maximal inhibitory concentration (IC50)** | purified MHC/competitive/fluorescence,  cellular MHC/competitive/fluorescence,  cellular MHC/competitive/radioactivity,  purified MHC/competitive/radioactivity,  cellular MHC/T cell inhibition,  binding assay |
| **off rate** | lysate MHC/direct/radioactivity,  cellular MHC/direct/fluorescence,  purified MHC/direct/fluorescence,  binding assay |
| **on rate** | cellular MHC/direct/fluorescence,  purified MHC/direct/fluorescence,  binding assay |
| **qualitative binding** | lysate MHC/direct/radioactivity,  purified MHC/direct/radioactivity,  purified MHC,  lysate MHC/direct/fluorescence,  purified MHC/competitive/fluorescence,  cellular MHC/competitive/fluorescence,  cellular MHC/competitive/radioactivity,  purified MHC/competitive/radioactivity,  purified MHC/direct/phage display,  lysate MHC,  cellular MHC,  purified MHC/direct/fluorescence,  cellular MHC/direct/fluorescence,  cellular MHC/T cell inhibition,c  ellular MHC/direct/radioactivity,  binding assay |

**Table S1:** valid Assay Responses measured in MHC binding assays and their associated valid Assay Types.

| Entity | Column | Description | Valid values |
| --- | --- | --- | --- |
| Peptide | Object type | Refers to whether a peptide is derived from a natural protein or synthetic. | Peptide from Protein **or** Peptide, No Natural Source |
| Peptide | Epitope Structure Definition | Refers to whether the Epitope Linear Sequence constitutes the exact sequence recognized by the MHC molecule or is a longer protein sequence containing the exact epitope | Exact Epitope **or** Epitope-containing region/antigenic site |

**Table S2:** descriptions of two additional peptide-specific fields that we tested (**Figure S1**).

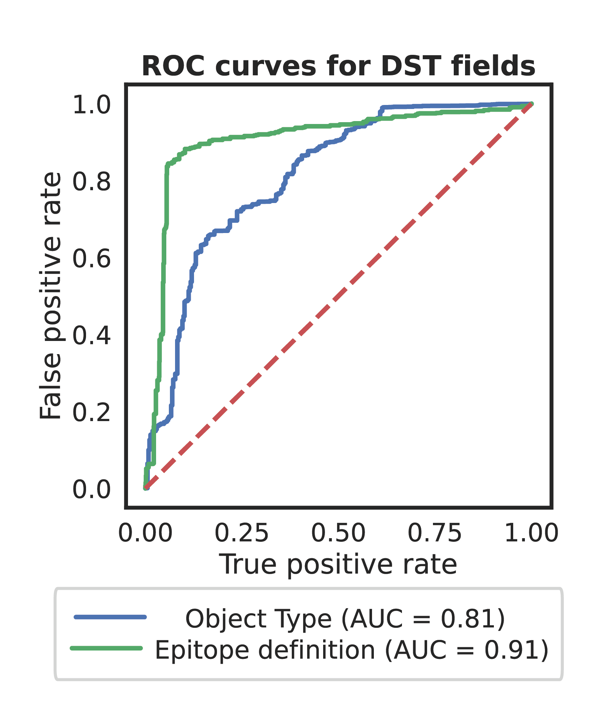

**Figure S1:**“Epitope Object Type” indicates whether a peptide is derived from a natural protein (“Peptide from Protein”) or is either a derivative of a non-natural protein or synthetic derivative of a peptide from a natural protein (“Peptide, no natural source”). The majority of peptides in the IEDB are derived from natural proteins; as a result, this is a highly imbalanced class (92.4% in the majority class). The VLM achieved a ROC-AUC of 0.81 and ROC-AUC_0.1 of 0.58. “Epitope Definition” refers to whether the peptide sequence is expected to be the exact epitope presented by the MHC molecule (“Exact Epitope”) or is contained within a longer sequence (“Epitope containing region/antigenic site”). This is calculated based on the peptide length and the classes of MHC molecule that the peptide is tested against within the paper. This exhibits a weaker imbalance in our dataset than Object Type (70.2% of peptides belong to the “Exact Epitope” class). The ROC-AUC was 0.91 and AUC_0.1 was 0.75.

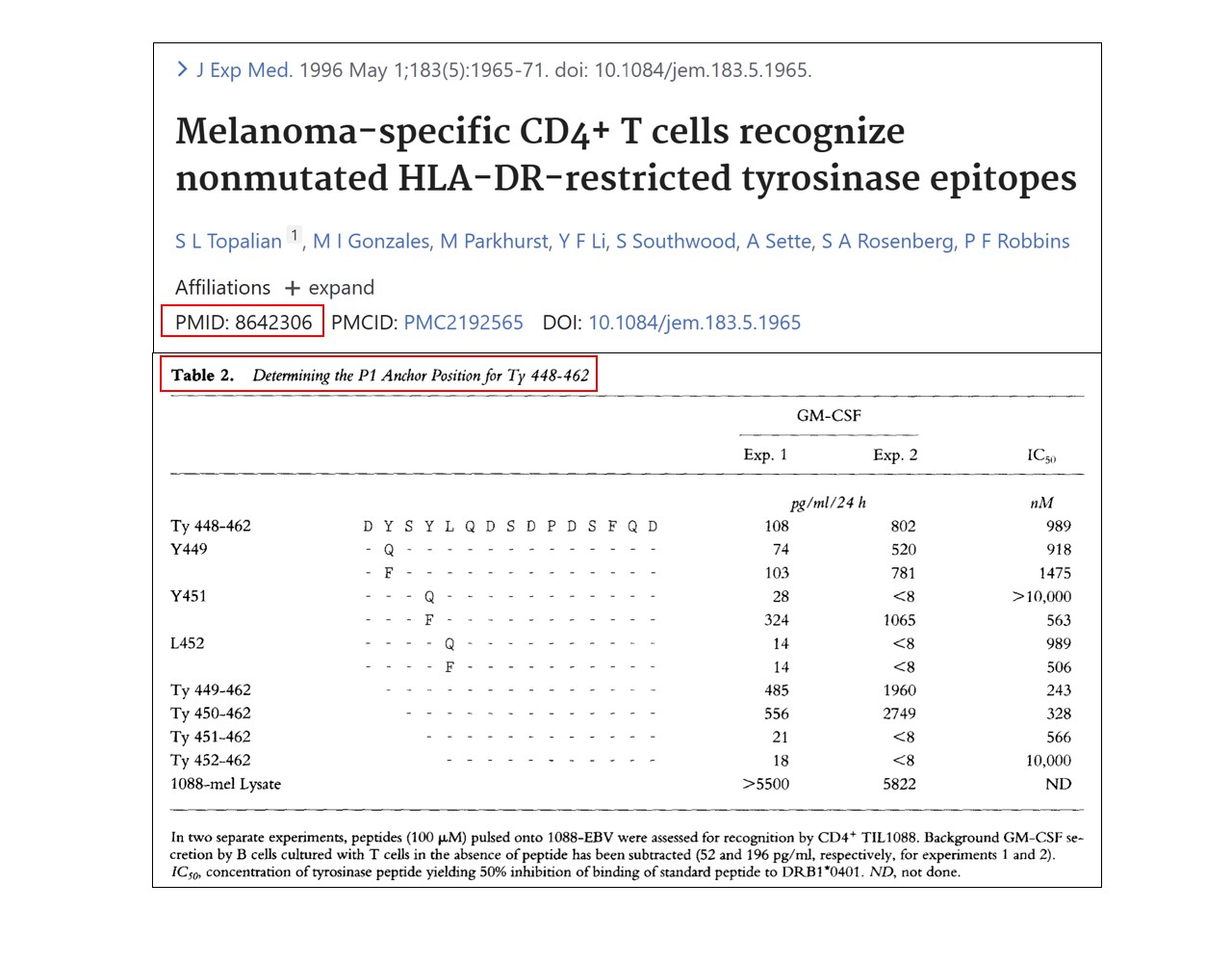

**Figure S2.** Given this table, the model sometimes determines GM-CSF as the assay response despite being provided with a fixed vocabulary, which includes the correct value, half maximal inhibitory concentration (IC50). Table reproduced from Topalian et al, 1996 (PMID: 8642306) with permission from the author Dr. Alessandro Sette.
